# ERGA-BGE genome of *Cheirolophus tagananensis*: an IUCN endangered shrub endemic to the Canary Islands

**DOI:** 10.1101/2025.01.24.634661

**Authors:** Jaume Pellicer, Teresa Garnatje, Daniel Vitales, Oriane Hidalgo, Joan Vallès, Alfredo García-Fernández, Arnoldo Santos-Guerra, Astrid Böhne, Rita Monteiro, Rosa Fernández, Nuria Escudero, Wellcome Sanger Institute Tree of Life Management, Samples and Laboratory Team, Wellcome Sanger Institute Scientific Operations, Sequencing Operations, Wellcome Sanger Institute Tree of Life Core Informatics Team, Abitha Thomas, Benjamin Jackson, Jonathan MD Wood, Kerstin Howe, Mark Blaxter, Shane McCarthy, Vianey Paola Barrera Enriquez, Leanne Haggerty, Fergal Martin, Chiara Bortoluzzi

## Abstract

The reference genome of *Cheirolophus tagananensis*, locally known as the Cabezón de Taganana, will provide an exceptional opportunity to establish a new framework to develop comparative genomic tools. These tools will help uncover the genetic basis of rapid plant radiations and microevolutionary adaptation processes of insular species on oceanic islands. This genomic resource will also contribute to facilitate the establishment of better informed *in situ* and *ex-situ* conservation strategies for this narrow endemic in the face of potential habitat degradation, and support taxonomic studies to better understand genetic diversity at the population, species, and genus levels. A total of 16 contiguous chromosomal pseudomolecules were assembled from the genome sequence. This chromosome-level assembly encompasses 0.62 Gb, composed of 421 contigs and 235 scaffolds, with contig and scaffold N50 values of 4.0 Mb and 36.5 Mb, respectively.

## Introduction

*Cheirolophus tagananensis* (Svent.) Holub, locally known as the Cabezón de Taganana, is a medium-sized woody shrub in the sunflower family, namely the *Asteraceae*, which produces multiple artichoke-like globose inflorescences with yellowish florets (Bramwell & Bramwell, 1974). The genus represents one of the most iconic and fastest plant radiations found in Macaronesia (Vitales et al., 2014), which gave rise to a significant number of micro endemisms. This shrub occurs exclusively on Tenerife (Canary Islands). Only a handful of populations can be found in the northernmost part of the island, across the Anaga Mountain range. This species grows primarily on rocky coastal cliffs, and exposed to ocean winds, where it is adapted to thrive in open areas together with other thermophilous taxa (Marrero et al., 2004). The chromosome number of *C. tagananensis* is unknown. However, most likely the species has a diploid chromosome number of 30-32 chromosomes, compatible with a paleopolyploid origin, based on an available karyological and genome size survey that includes the analysis of closely related species from the archipelago (Hidalgo et al., 2017).

*Cheirolophus tagananensis* is currently classified as ‘Vulnerable’ according to the IUCN Red List of Threatened Species (https://www.iucnredlist.org/species/165120/5975940) and ‘Endangered’ based on the Spanish Red List of Threatened Flora (Marrero et al., 2004). The primary reason for such conservation status is underpinned by the restricted distribution of this species. This is only confirmed to a reduced area of 3 km^2^ in isolated populations. In addition, according to the Centinela Database for the Government of the Canary Islands (https://www.biodiversidadcanarias.es/centinela/especie/F01523), *C. tagananensis* is also under protection (Appendix I) of the Bern Convention given its relevance for the preservation of Canarian ecosystems and habitats included in the Directive 92/43/CEE. Fully developed specimens of *C. tagananensis* present multiple inflorescence clusters, which serve as a food source for a variety of insects, thus contributing to the functioning of island ecosystems. Immature seeds are often predated by insect larvae, which compromise their propagation and fitness (Gómez Campo, 1996).

The generation of this reference resource was coordinated by the European Reference Genome Atlas (ERGA) initiative’s Biodiversity Genomics Europe (BGE) project, supporting ERGA’s aims of promoting transnational cooperation to promote advances in the application of genomics technologies to protect and restore biodiversity (Mazzoni et al., 2023).

## Materials & Methods

ERGA’s sequencing strategy includes Oxford Nanopore Technology (ONT) and/or Pacific Biosciences (PacBio) for long-read sequencing, along with Hi-C sequencing for chromosomal architecture, Illumina Paired-End (PE) for polishing (i.e. recommended for ONT-only assemblies), and RNA sequencing for transcriptome profiling, to facilitate genome assembly and annotation.

### Sample and Sampling Information

On 22 September 2023, fresh leaves from one adult living plant cultivated in the greenhouse facility of the Botanical Institute of Barcelona (Spain) were collected by Jaume Pellicer, placed in liquid nitrogen, and stored at -80°C until DNA extraction. Plants in the greenhouse facility in Barcelona were kept in cultivation from a previous field expedition conducted in 2014 by Daniel Vitales, Arnoldo Santos-Guerra, and Alfredo García under the permit number AFF 265/14, issued by the Cabildo Insular de Tenerife, Area de Medio Ambiente, Sostenibilidad Territorial y de Recursos y Aguas, Gobierno de Canarias, which granted permission to collect a reduced number of seeds from 10 adult plants (monoecious hermaphrodite) of *Cheirolophus tagananensis*. The species was identified by specialist local botanist Arnoldo Santos-Guerra using the identification key for the genus in “Libro rojo de especies vegetales amenazadas de las Islas Canarias” by Gómez Campo (1996).

### Vouchering information

A physical herbarium voucher of the specimen sequenced is deposited at Herbarium MA (Real Jardín Botánico de Madrid, CSIC), https://rjb.csic.es/rjb-colecciones/herbario-ma/ under the herbarium voucher ID MA-01-00963497.

Frozen reference tissue and DNA samples from the same individual are deposited at the Biobank of the Museo Nacional de Ciencias Naturales (MNCN, CSIC) https://mncn.csic.es/en under voucher ID MCN-ADN-151769/71. Also, seeds from other individuals than the sequenced ones have been deposited at Rey Juan Carlos University Germplasm bank https://bgurjc.weebly.com/ with the reference number BG URJC:273 - 1.

### Data Availability

*Cheirolophus tagananensis* and the related genomic study were assigned to Tree of Life ID (ToLID) ‘daCheTaga1’ and all sample, sequence, and assembly information are available in ENA under the umbrella BioProject PRJEB76612. The sample information is available at the following BioSample accession: SAMEA114757429. The genome assembly is accessible from ENA under accession number GCA_964300405.1 and the annotated genome is available through the Ensembl Beta website (https://projects.ensembl.org/erga-bge/).

Sequencing data produced as part of this project are available from ENA at the following accessions: ERX13020692 and ERX12671966. Documentation related to the genome assembly and curation can be found in the ERGA Assembly Report (EAR) document available at https://github.com/ERGA-consortium/EARs/blob/main/Assembly_Reports/Cheirolophus_tagananensis/daCheTga1. Further details and data about the project are hosted on the ERGA portal at https://portal.erga-biodiversity.eu/data_portal/65001.

### Genetic Information

The estimated genome size of *C. tagananensis* is between 0.67 Gb (Garnatje et al., 2007) and 0.69 Gb (Hidalgo et al., 2017). This is a diploid genome with a haploid number of 15-16 chromosomes (2n = 30-32) based on a study conducted across the genus. All information for this species was retrieved from Genomes on a Tree (Challis et al., 2023).

### DNA/RNA processing

The workflow for high molecular weight (HMW) DNA extraction at the Wellcome Sanger Institute (WSI) Tree of Life Core Laboratory includes a sequence of procedures: sample preparation and homogenisation, DNA extraction, fragmentation, and purification. Detailed protocols are available on protocols.io (Denton et al., 2023). The daCheTaga1 sample was prepared for DNA extraction by weighing and dissecting it on dry ice (Jay et al., 2023). Tissue from the leaf tissue was homogenised by cryogenic bead beating (Jackson & Howard, 2023a). HMW DNA was extracted using the Plant Organic Extraction protocol (Jackson & Howard, 2023b). HMW DNA was sheared using the Covaris g-Tube protocol (Sampaio et al., 2023). Sheared DNA was purified by solid-phase reversible immobilisation, using AMPure PB beads to eliminate shorter fragments and concentrate the DNA (Sampaio & Howard, 2023). The concentration of the sheared and purified DNA was assessed using a Nanodrop spectrophotometer and Qubit Fluorometer using the Qubit dsDNA High Sensitivity Assay kit. Fragment size distribution was evaluated by running the sample on the FemtoPulse system.

Hi-C data were generated from leaf tissue from the daCheTaga1 using the Arima-HiC v2 kit. Tissue was finely ground using cryoPREP and then subjected to nuclei isolation. Nuclei were isolated using a modified protocol of the Qiagen QProteome Cell Compartment Kit where only CE1 and CE2 buffers are used in combination with QiaShredder spin columns. After isolation, the nuclei were fixed using 37% formaldehyde solution to crosslink the DNA. The crosslinked DNA was then digested using the restriction enzyme master mix. The 5’-overhangs were then filled in and labelled with biotinylated nucleotides and proximally ligated. An overnight incubation was carried out for enzymes to digest the remaining proteins and for crosslinks to reverse. A clean-up was performed with SPRIselect beads before library preparation. DNA concentration was quantified using the Qubit Fluorometer v2.0 and Qubit HS Assay Kit according to the manufacturer’s instructions.

RNA Extraction: Automated MagMax™ mirVana protocol (do Amaral et al., 2023). The RNA concentration was assessed using a Nanodrop spectrophotometer and a Qubit Fluorometer using the Qubit RNA Broad-Range Assay kit. Analysis of the integrity of the RNA was done using the Agilent RNA 6000 Pico Kit and Eukaryotic Total RNA assay.

### Library Preparation and Sequencing

Library preparation and sequencing were performed at the WSI Scientific Operations core. At the minimum, HMW DNA samples were required to have an average fragment size exceeding 8 kb and a total mass over 400 ng to proceed to the low input SMRTbell Prep Kit 3.0 protocol (PacBio, California, USA), depending on genome size and sequencing depth required. Libraries were prepared using the SMRTbell Prep Kit 3.0 (Pacific Biosciences, California, USA) as per the manufacturer’s instructions. The kit includes the reagents required for end repair/A-tailing, adapter ligation, post-ligation SMRTbell bead cleanup, and nuclease treatment. Following the manufacturer’s instructions, size selection and clean-up were carried out using diluted AMPure PB beads (PacBio, California, USA). DNA concentration was quantified using the Qubit Fluorometer v4.0 (Thermo Fisher Scientific) with Qubit 1X dsDNA HS assay kit and the final library fragment size analysis was carried out using the Agilent Femto Pulse Automated Pulsed Field CE Instrument (Agilent Technologies) and gDNA 55kb BAC analysis kit.

Prepared libraries were normalised to 2nM and 15μL used for making complexes. For libraries below 2nM all 10uL was used for making complexes. Primers were annealed and polymerases were hybridised to create circularised complexes according to the manufacturer’s instructions. The complexes were purified with the 1.2X clean-up with SMRTbell beads. The purified complexes were then diluted to the Revio loading concentration, between 200 - 300pM, and spiked with a Revio sequencing internal control.

Samples were sequenced using the Revio system on Revio 25M SMRT cells (PacBio, California, USA). The SMRT link software, a PacBio web-based end-to-end workflow manager, was used to set up and monitor the run, as well as perform primary and secondary analysis of the data upon completion.

For Hi-C library preparation, DNA was fragmented to a size of 400 to 600 bp using a Covaris E220 sonicator. The DNA was then enriched, barcoded, and amplified using the NEBNext Ultra II DNA Library Prep Kit (New England Biolabs) following the manufacturer’s instructions. The Hi-C sequencing was performed using paired-end sequencing with a read length of 150 bp on an Illumina NovaSeq X instrument.

Poly(A) RNA-Seq libraries were constructed using the NEB Ultra II RNA Library Prep kit, following the manufacturer’s instructions. RNA sequencing was performed on the Illumina NovaSeq X instrument.

In total, 94x PacBio and 154x HiC data were sequenced to generate the assembly.

### Genome Assembly Methods

The HiFi reads were assembled using Hifiasm (Cheng et al., 2021) in Hi-C phasing mode, where data were separated into two haplotypes. These haplotypes were then curated to generate a final assembly. The Hi-C reads were aligned to the contigs using bwa-mem2 (Vasimuddin et al., 2019), and contigs were scaffolded with YaHS (Zhou et al., 2023), using the --break option for handling potential misassemblies. The resulting scaffolded assemblies were evaluated using Gfastats (Formenti et al., 2022), BUSCO (Manni et al., 2021), and MERQURY.FK (Rhie et al., 2020).

Both mitochondrial and plastid genomes were assembled using oatk (Zhou et al., 2024).

The assembly was decontaminated using the Assembly Screen for Cobionts and Contaminants (ASCC) pipeline (article in preparation). Flat files and maps used in curation were generated in TreeVal (Pointon et al., 2023). Manual curation was primarily conducted using PretextView (Harry, 2022), with additional insights provided by JBrowse2 (Diesh et al., 2023) and HiGlass (Kerpedjiev et al., 2018). Scaffolds were visually inspected and corrected as described by (Howe et al., 2021). Any identified contamination, missed joins, and mis-joins were corrected, and duplicate sequences were tagged and removed. The curation process is documented at https://gitlab.com/wtsi-grit/rapid-curation (article in preparation). Summary analysis of the released assembly was performed using the ERGA-BGE Genome Report ASM Galaxy workflow (10.48546/workflowhub.workflow.1104.1).

### Genome Annotation Methods

A gene set was generated using the Ensembl Gene Annotation system (Aken et al., 2016), primarily by aligning publicly available short-read RNA-seq data from BioSample: SAMEA114757429 to the genome. Gaps in the annotation were filled via protein-to-genome alignments of a select set of clade-specific proteins from UniProt (Consortium, 2019), which had experimental evidence at the protein or transcript level. At each locus, data were aggregated and consolidated, prioritising models derived from RNA-seq data, resulting in a final set of gene models and associated non-redundant transcript sets. To distinguish true isoforms from fragments, the likelihood of each open reading frame (ORF) was evaluated against known metazoan proteins. Low-quality transcript models, such as those showing evidence of fragmented ORFs, were removed.

In cases where RNA-seq data were fragmented or absent, homology data were prioritised, favouring longer transcripts with strong intron support from short-read data. The resulting gene models were classified into two categories: protein-coding, and long non-coding. Models that did not overlap protein-coding genes, and were constructed from transcriptomic data were considered potential lncRNAs. Potential lncRNAs were further filtered to remove single-exon loci due to their unreliability. Putative miRNAs were predicted by performing a BLAST search of miRBase (Kozomara et al., 2019) against the genome, followed by RNAfold analysis (Gruber et al., 2008). Other small non-coding loci were identified by scanning the genome with Rfam (Kalvari et al., 2018) and passing the results through Infernal (Nawrocki & Eddy, 2013). Summary analysis of the released annotation was carried out using the ERGA-BGE Genome Report ANNOT Galaxy workflow (10.48546/workflowhub.workflow.1096.1).

## Results

### Genome Assembly

The genome assembly has a total length of 624,001,908 bp with 92.1 % of the sequence assigned to 16 chromosomes plus the mitochondrial genome and plastid (chloroplast genome) (Figures 1 and 2), with an overall GC content of 36.6%. It has a contig N50 of 4.0 Gb (L50 = 50) and a scaffold N50 of 36.5 Gb (L50 = 7). There are 186 gaps, totalling 37.2 kb in cumulative size. The single-copy gene content analysis using the Eudicots database with BUSCO (Manni et al., 2021) resulted in 98.0% completeness (92.7% single and 5.3% duplicated; for an interactive plot, refer to the BlobToolkit viewer). 93.2% of reads k-mers were present in the assembly and the assembly has a base accuracy Quality Value (QV) of 62.0 as calculated by Merqury (Rhie et al., 2020).

**Figure 1.**
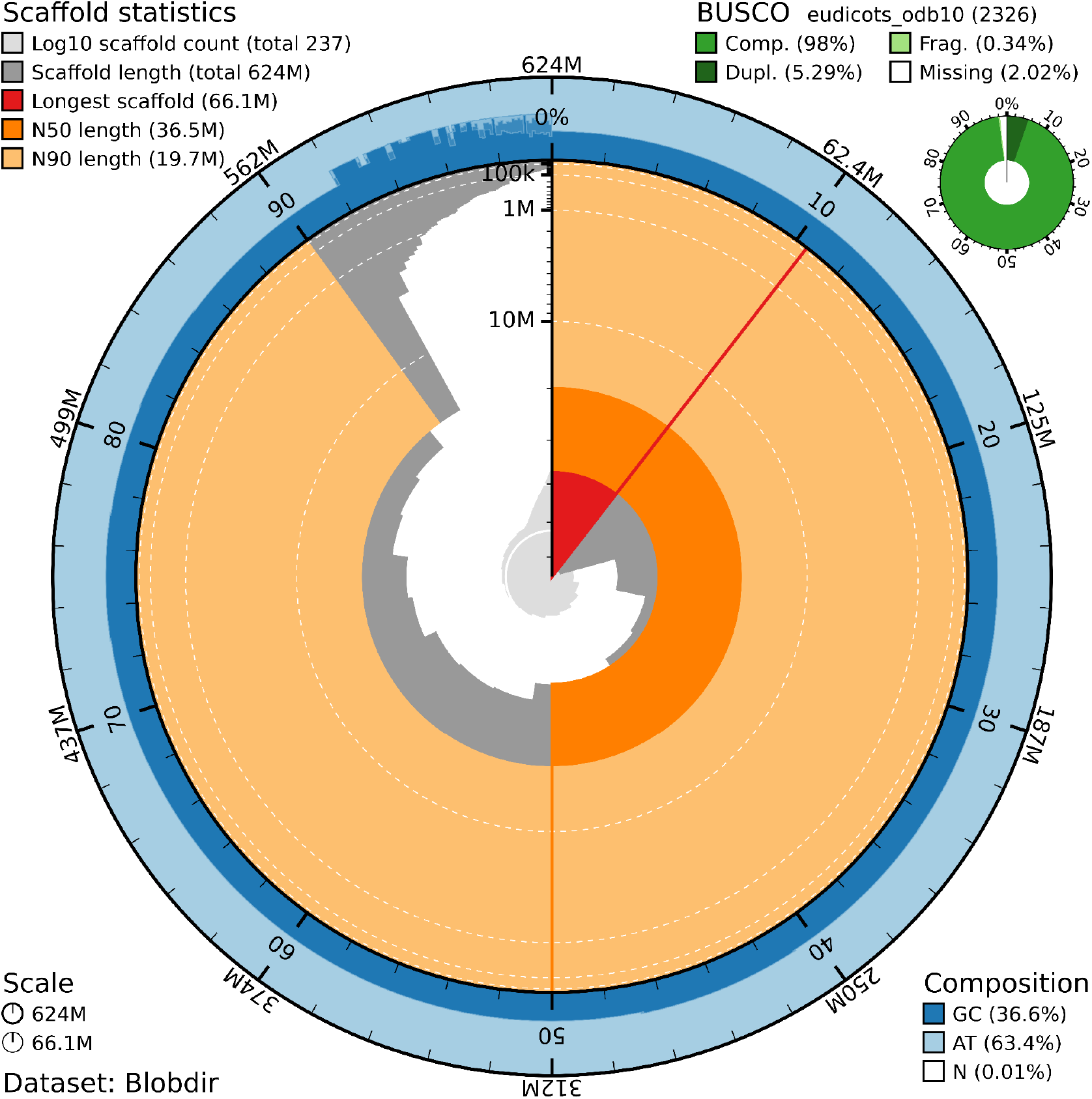
Snail plot summary of assembly statistics. The main plot is divided into 1,000 size-ordered bins around the circumference, with each bin representing 0.1% of the 624,001,908 bp assembly including the mitochondrial genome. The distribution of sequence lengths is shown in dark grey, with the plot radius scaled to the longest sequence present in the assembly (66.1 Mb, shown in red). Orange and pale-orange arcs show the scaffold N50 and N90 sequence lengths (36,486,517 and 19,666,334 bp), respectively. The pale grey spiral shows the cumulative sequence count on a log-scale, with white scale lines showing successive orders of magnitude. The blue and pale-blue area around the outside of the plot shows the distribution of GC, AT, and N percentages in the same bins as the inner plot. A summary of complete, fragmented, duplicated, and missing BUSCO genes found in the assembled genome from the Eudicots database (odb10) is shown on the top right.

**Figure 2.**
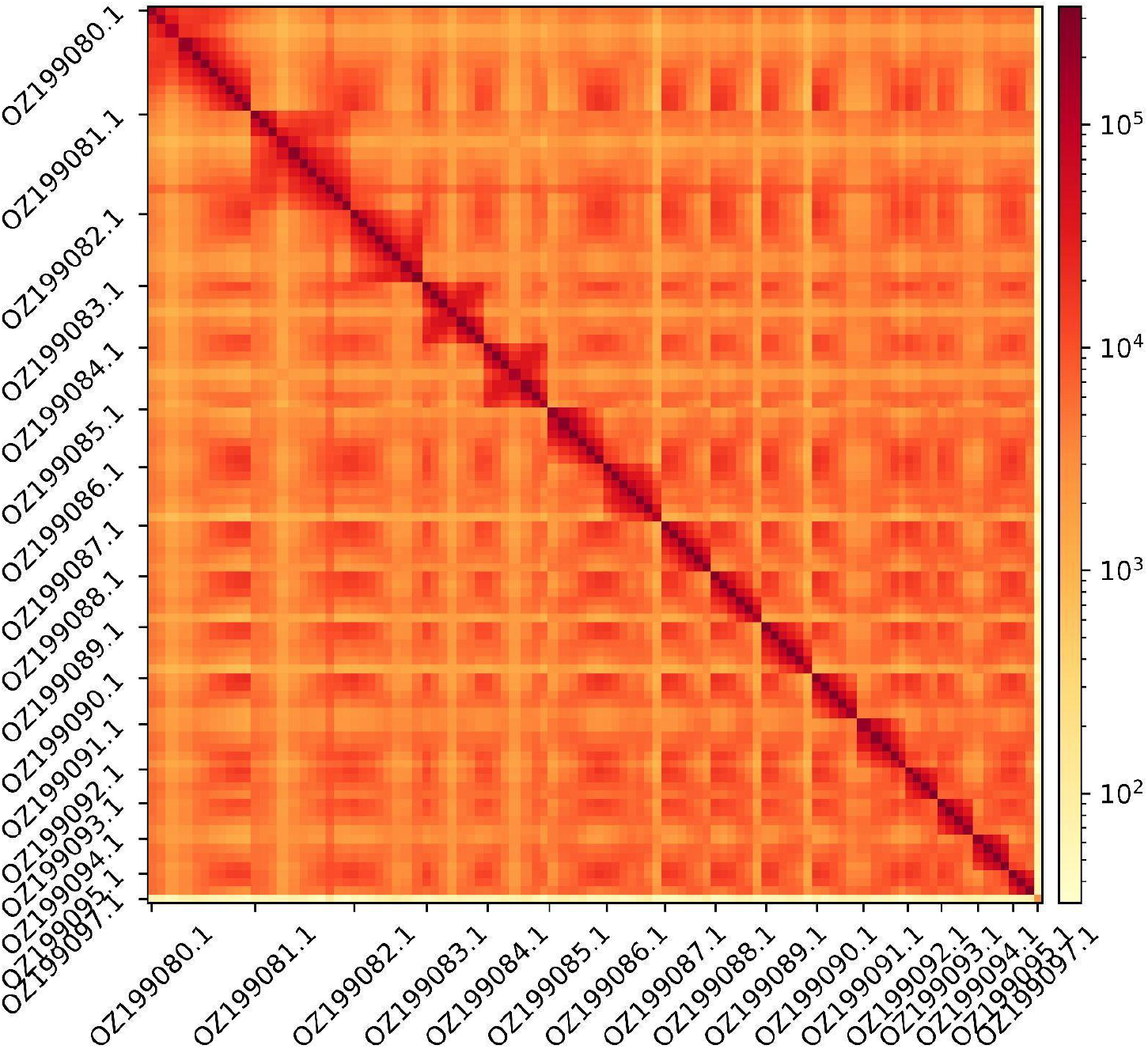
Hi-C contact map showing spatial interactions between regions of the genome. The diagonal corresponds to intra-chromosomal contacts, depicting chromosome boundaries. The frequency of contacts is shown on a logarithmic heatmap scale. Hi-C matrix bins were merged into a 100 kb bin size for plotting. The plastid genome is identified by GenBank accession OZ199097.1.

### Genome Annotation

The genome annotation consists of 19,080 protein-coding genes with associated 22,290 transcripts, in addition to 19,488 non-coding genes (Table 1). Using the longest isoform per transcript, the single-copy gene content analysis using the Eudicots odb10 database with BUSCO resulted in 79.2% completeness. Using the OMAmer Viridiplantae-v2.0.0.h5 database for OMArk (Nevers et al., 2025) resulted in 82.9% completeness and 94.9% consistency (Table 2).

**Table 1.** Statistics from assembled gene models.

|  | No. genes | No. transcripts | Mean gene length (bp) | No. single-exon genes | Mean exons per transcript |
| --- | --- | --- | --- | --- | --- |
| mRNA | 19,080 | 22,290 | 5,325 | 1,373 | 6.5 |
| pseudogene | 0.0 | 0.0 | 0.0 | 0.0 | 0.0 |
| snoRNA | 680 | 680 | 105 | 680 | 1.0 |
| lncRNA | 1,379 | 1,595 | 2,554 | 5 | 2.6 |
| miRNA | 0.0 | 0.0 | 0.0 | 0.0 | 0.0 |
| snRNA | 204 | 204 | 149 | 204 | 1.0 |
| rRNA | 16,314 | 16,314 | 1,581 | 16,314 | 1.0 |
| scRNA | 0.0 | 0.0 | 0.0 | 0.0 | 0.0 |
| Other ncRNA | 911 | 911 | 74-133 | 911 | 1.0-1.0 |

**Table 2.** Annotation completeness and consistency scores calculated by BUSCO run in protein mode (eudicots_odb10) and OMArk (Viridiplantae-v2.0.0.h5)

|  | Complete | Single copy | Duplicated | Fragmented | Missing |
| --- | --- | --- | --- | --- | --- |
| BUSCO | 1,844 (79.2%) | 1,743 (74.9%) | 101 (4.3%) | 124 (5.3%) | 358 (15.5%) |
| OMArk | 9,287 (82.9%) | 7,066 (63.1%) | 2,221 (19.8%) | - | 1,911 (17.0%) |
|  | Consistent | Inconsistent | Contaminants | Unknown |  |
| OMArk | 18,122 (94.9%) | 253 (1.3%) | 0.0 (0.0%) | 705 (3.7%) |  |

## Acknowledgements

We would like to thank the Cabildo Insular de Tenerife (Gobierno de Canarias) for their support in issuing collecting permits. Fieldwork and sample collection were also possible thanks to the support of the Linnean Society and the Systematic Foundation under the “Unraveling the *Cheirolophus webbianus* complex in the north of Tenerife Island” project. Miquel Veny is also thanked for his assistance and maintenance of specimens in cultivation at the facilities of the Botanical Institute of Barcelona. We acknowledge the support of the Freiburg Galaxy Team: Saim Momin and Björn Grüning, Bioinformatics, University of Freiburg (Germany), funded by the German Federal Ministry of Education and Research BMBF grant 031 A538A de.NBI-RBC and the Ministry of Science, Research and the Arts Baden-Württemberg (MWK) within the framework of LIBIS/de.NBI Freiburg. We would like to acknowledge the assembly reviewer, Lola Demirdjian from Genoscope.

## Author Contributions

JP coordinated the project; JP, DV, AF and ASG collected the species; ASG identified the species; JP sampled and preserved biological material and provided metadata; AsB, NE, RF and RM provided support in sampling, shipping of biological material, metadata collection, and management; AT, BJ, and the WSI ToL M, S and LT extracted DNA and prepared libraries, WSI ToL SO performed sequencing; JMDW and WSI ToL IT performed genome assembly and curation under the supervision of KH, MB, and SMcC; LH, SS, and FM performed genome annotation; CB generated the analysis and report. All authors contributed to the writing, review, and editing of this genome note and read and approved the final version.

## Author Information

Members of the Wellcome Sanger Institute Tree of Life Management, Samples and Laboratory team are listed here: https://doi.org/10.5281/zenodo.12162482.

Members of Wellcome Sanger Institute Scientific Operations: Sequencing Operations are listed here: https://doi.org/10.5281/zenodo.12165051.

Members of the Wellcome Sanger Institute Tree of Life Core Informatics team are listed here: https://doi.org/10.5281/zenodo.12160324.

